# CRISPR/Cas9 cleavages in budding yeast reveal templated insertions and strand-specific insertion/deletion profiles

**DOI:** 10.1101/193466

**Authors:** Brenda R. Lemos, Adam C. Kaplan, Ji Eun Bae, Alex Ferazzoli, James Kuo, Ranjith P. Anand, David P. Waterman, James E. Haber

**Affiliations:** Department of Biology and Rosenstiel Basic Medical Sciences Research Center, Brandeis University, Waltham, MA 02454, USA

**Author notes:** These authors contributed equally to this work (Brenda R. Lemos, Adam C. Kaplan).

## Abstract

Harnessing CRISPR-Cas9 technology has provided an unprecedented ability to modify genomic loci via DNA double-strand break (DSB) induction and repair. We have analyzed nonhomologous end-joining (NHEJ) repair induced by Cas9 in the budding yeast *Saccharomyces cerevisiae* and find that the orientation of binding of Cas9 and its guide RNA (gRNA) profoundly influences the pattern of insertion/deletions (indels) at the site of cleavage. A common indel created by Cas9 is a one base pair (+1) insertion that appears to result from Cas9 creating a 1-bp 5’ overhang that is filled in by a DNA polymerase and ligated. The origin of +1 insertions was investigated by using two gRNAs with PAM sequences located on opposite DNA strands but designed to cleave the same sequence. These templated +1 insertions are dependent on the X-family DNA polymerase, Pol4. Deleting Pol4 also eliminated +2 and +3 insertions, which were biased toward homonucleotide insertions. Using inverted PAM (iPAM) sequences, we also found significant differences in overall NHEJ efficiency and repair profiles, suggesting that the binding of the Cas9::gRNA complex influences subsequent NHEJ processing. As with well-studied events induced by the site-specific HO endonuclease, CRISPR-Cas9 mediated NHEJ repair depends on the Ku heterodimer and DNA ligase 4. Cas9 events, however, are highly dependent on the Mre11-Rad50-Xrs2 complex, independent of Mre11’s nuclease activity. Inspection of the outcomes of a large number of Cas9 cleavage events in mammalian cells (van Overbeek et al., 2016) reveals a similar templated origin of +1 insertions in human cells, but also a significant frequency of similarly templated +2 insertions.

## INTRODUCTION

The rapid emergence of tools to exploit the CRISPR-Cas9 RNA guided endonuclease has accelerated genome editing in systems ranging from yeast to mammals (DiCarlo et al., 2013; Mali et al., 2013; Shen et al., 2013). Guided endonucleases target site-specific DNA to create double-strand breaks (DSBs) (Hsu et al., 2014), which can be repaired either by non-homologous end joining (NHEJ) or homologous recombination (HR) (reviewed by (Mehta and Haber, 2014)). The mechanisms of these repair processes, including the core genetic components required, have been extensively studied *in vivo* using the site-specific HO endonuclease of *Saccharomyces cerevisiae* (Haber, 2016). We were therefore interested in comparing the DSB repair induced by CRISPR-Cas9 with events promoted by HO endonuclease in budding yeast.

The *Streptococcus pyogenes* Cas9 endonuclease scans the genome to locate Protospacer Adjacent Motifs (PAM) of 5’ NGG 3’, which are required for binding and cleavage. Specificity is achieved by a Cas9-bound guide RNA (gRNA) that pairs with sequences adjacent to the PAM, resulting in double-strand DNA cleavage 3-bp 5’ of the PAM sequence (Doudna and Charpentier, 2014). In contrast, HO endonuclease recognizes a defined 24-bp cleavage site located in the *S. cerevisiae* mating type (*MAT*) locus. When placed under control of a galactose-inducible promoter, HO cleaves euchromatic targets within 30-60 min, creating 4-nt 3’ overhanging ends (Haber, 2002; Hicks et al., 2011; Nickoloff et al., 1986).

NHEJ can result either in faithful re-ligation of the broken ends, or in the mutagenic insertions or deletions of base pairs at the junctions (indel formation). In budding yeast, perfect re-ligation of the 3’ overhanging DSB ends requires the “classic” evolutionarily conserved end-joining machinery including the Ku70-Ku80 proteins as well as DNA ligase 4 and its associated Lif1 (Xrcc4 homolog) and Nej1 co-factors (Moore and Haber, 1996; Valencia et al., 2001). Continuous expression of HO eliminates precise re-joinings by re-cutting, thereby forcing the recovery of mutated HO recognition sequences, including short, templated fill-ins and 3-bp deletions, all of which result from offset base pairings and subsequent processing of overhanging ends (Moore and Haber, 1996). The fill-ins depend on the Mre11-Rad50-Xrs2 (MRX) complex as well as the Polx family DNA polymerase, Pol4, whereas 3-bp deletions proved to be MRX and Pol4-independent. Additional microhomology-mediated end-joining events, resulting in larger deletions, can also be recovered with the same genetic requirements; but there is also a Ku-independent microhomology-mediated end-joining (MMEJ) process (Ma et al., 2003).

The CRISPR-Cas9 system can be used to create targeted indels by designing gRNAs homologous to the genomic site of interest. Biochemical experiments have shown that Cas9 cleaves 3-bp, 5’ from the PAM, resulting in a DSB, with Cas9 remaining bound to its target for hours after cleavage (Gasiunas et al., 2012; Jinek et al., 2012). *In vitro*, the 3’ end of the DSB that is not complementary to the gRNA is released sooner, raising the possibility that there could be strong biases in DSB end-processing dictated by the Cas9 protein (Richardson et al., 2016).

We were especially interested in whether we could infer the consequences of persistent Cas9 binding to the target site by creating the same cleavage in two different ways, by designing gRNAs to PAMs located symmetrically on either strand. Using several pairs of inverted PAM sequences (iPAMs), we found frequent +1 insertions, in which the added base is dependent on the orientation of Cas9::gRNA binding. These insertions appear to have arisen from Cas9 cleavages resulting 1-nt 5’ overhanging ends that are filled in by DNA polymerase Pol4. Moreover, there are substantial differences in the overall patterns of indels depending on the strand to which Cas9 is paired and bound.

## MATERIAL AND METHODS

### Parental strains and selection

Strains JKM179 (*ho*Δ *MAT*α *hml*Δ*::ADE1 hmr*Δ*::ADE1 ade1-100 leu2-3,112 lys5 trp1::hisG*′ *ura3-52 ade3::GAL::HO)* and JKM139 (*ho*Δ *MATa hml*Δ*::ADE1 hmr*Δ*::ADE1 ade1-100 leu2-3,112 lys5 trp1::hisG*′ *ura3-52 ade3::GAL::HO)* (Lee et al., 1998) were used in experiments targeting the *MAT* locus. These strains lack the heterochromatic *HML* and *HMR* donor sequences that would allow repair of a DSB at *MAT* by gene conversion; hence all repair is through NHEJ. Parental strain BY4733 (*MATa his3Δ200 trp1Δ63 leu2Δ0 met15Δ0 ura3Δ0)* was used for *CAN1* and *LYS2* Cas9 targeting experiments. ORFs were deleted by replacing the target with a prototrophic or antibiotic-resistance markers via single-step transformation of *Saccharomyces cerevisiae* colonies with PCR fragments (Rothstein, 1983).

### Plasmid construction

Plasmid pJD1, carrying Cas9 driven by the *GAL1* promoter was provided by the Church lab (DiCarlo et al., 2013). Each gRNA targeting the *MAT* locus, with an upstream SNR52 promoter and SUP4 terminator 3’ sequence was synthesized as a gBlock by IDT Technologies© and was cloned into plasmid pRS426 (*URA3*) by gap repair. gBlocks were designed with 50bp flanking homology to pRS426, and recombined *in vivo* after co-transforming 0.02 pmole gRNA gBlock and 5μg BamHI digested pRS426 (Oldenburg et al., 1997).

gRNAs gCAN1-1^W^, gCAN1-1^C^, and gLYS(1-4) were cloned into a *LEU2*-marked plasmid (bRA77), where Cas9 is expressed under a galactose-inducible promoter. Briefly, gRNAs were ligated into a BplI digested site, cloned in *E. coli*. Plasmids were verified by sequencing (GENEWIZ) and transformed into our yeast strains. A more detailed description of the method is presented by (Anand et al., 2017).

### Galactose induction and viability

Cas9 plasmids were transformed into parental strains using the conventional lithium acetate protocol (Gietz and Schiestl, 2007). Experiments targeting the *CAN1* and *LYS2* locus used the parental strain BY4733. Plasmids containing a galactose-inducible Cas9 and a constitutively expressed gRNA were introduced by transformation using the conventional lithium acetate protocol (Gietz and Schiestl, 2007). Cells were plated on YPD plates, and replica plated onto leucine drop-out plates for transformants. Cells containing the plasmid were serially diluted from 100,000 cells to 100 cells. Cells were plated onto leucine-dropout plates, either containing galactose to induce Cas9 expression or containing dextrose as a control. Experiments were performed in duplicates and repeated at least three times. Can^r^ colonies were selected by replica plating survivors onto canavanine-containing plates. The proportion of Lys2^+^ and Lys2^-^ colonies were identified by replica-plating survivors onto lysine-dropout plates.

For assays targeting the *MAT* locus, cells were grown at 30 °C overnight in synthetic medium lacking uracil and leucine and 2% dextrose, refreshed in selective medium containing 2% raffinose for 6 h, and induced in selective medium containing 2% galactose. Viability was determined as the average number of colonies growing with galactose relative to those on dextrose plates after 3 days at 30 °C. Viability assays were performed with three biological replicates. Viabilities were corrected by discounting the small proportion of cells that did not contain a *matα*1 mutation. Viability of strains with an unrepaired HO endonuclease-induced DSB were determined as previously described (Moore and Haber, 1996).

### DNA Sequence analysis

Using flanking primers to the DNA region of interest, PCR was used to amplify DNA from survivor colonies. PCR products were purified and Sanger sequenced using GENEWIZ, Inc (Cambridge, MA) and Eton Bioscience, Inc (Boston, MA). Data represent samples from at least 3 independent inductions of Cas9 cleavage. Barcoded primers were used to amplify *LYS2* from a pool of either gLYS2-4^C^ or gLYS2-4^W^ survivors. Amplified PCR samples were purified and sequenced using MiSeq Illumina® sequencing technology.

## RESULTS

### NHEJ of Cas9 DSBs reveals frequent +1-bp templated insertions

In several reports of Cas9-generated indels in mammalian cells, there are frequent instances of +1 insertions (Jiang et al., 2014; Seeger and Sohn, 2016; van Overbeek et al., 2016). Close inspection of these sequences suggested that the insertions were nonrandom and in fact could be explained if Cas9 sometimes leaves a 1-nt 5’ overhang by cleaving 3-nt from the PAM on the gRNA complementary strand and 4-nt away from the PAM on the non-complementary strand that would then be filled in and ligated (Fig. 1B and 1C). We found that this was also the case in budding yeast. One example is shown in Fig. 1B, using a gRNA (LYS2-2^C^) that cleaves within the *LYS2* locus. In these experiments, a *LEU2*-marked plasmid carrying both a gRNA expressed constitutively and a galactose-inducible Cas9 was transformed into strain BY4733 and then cells plated on synthetic medium lacking leucine and with galactose as the carbon source (Anand et al., 2017). Viability was approximately 1%, of which about 75% of the viable colonies were Lys2^-^ (Fig. 1F). Among the Lys2^-^ survivors (Fig. 1D), 35% had the insertion of a single T, the nucleotide expected if there were a 1-nt 5’ overhang after Cas9 cleavage. At substantially lower frequencies there were +1 insertions of G, C and A at the same site. Another 15% had a single base-pair deletion. The Lys2^+^ survivors all proved to have in-frame deletions (3 or 6-bp) or insertions (3-bp), or else were base pair substitutions (Figure 1D). Among the Lys2^+^ events, the most prominent was a 3-bp deletion that would arise through single-strand annealing involving TTG microhomologies flanking a blunt-ended cut; such deletions would arise after 5’ to 3’ resection of the ends and annealing of the single-stranded, 3’ ended AAC and TTG sequences at the two DSB ends.

**Figure 1.**
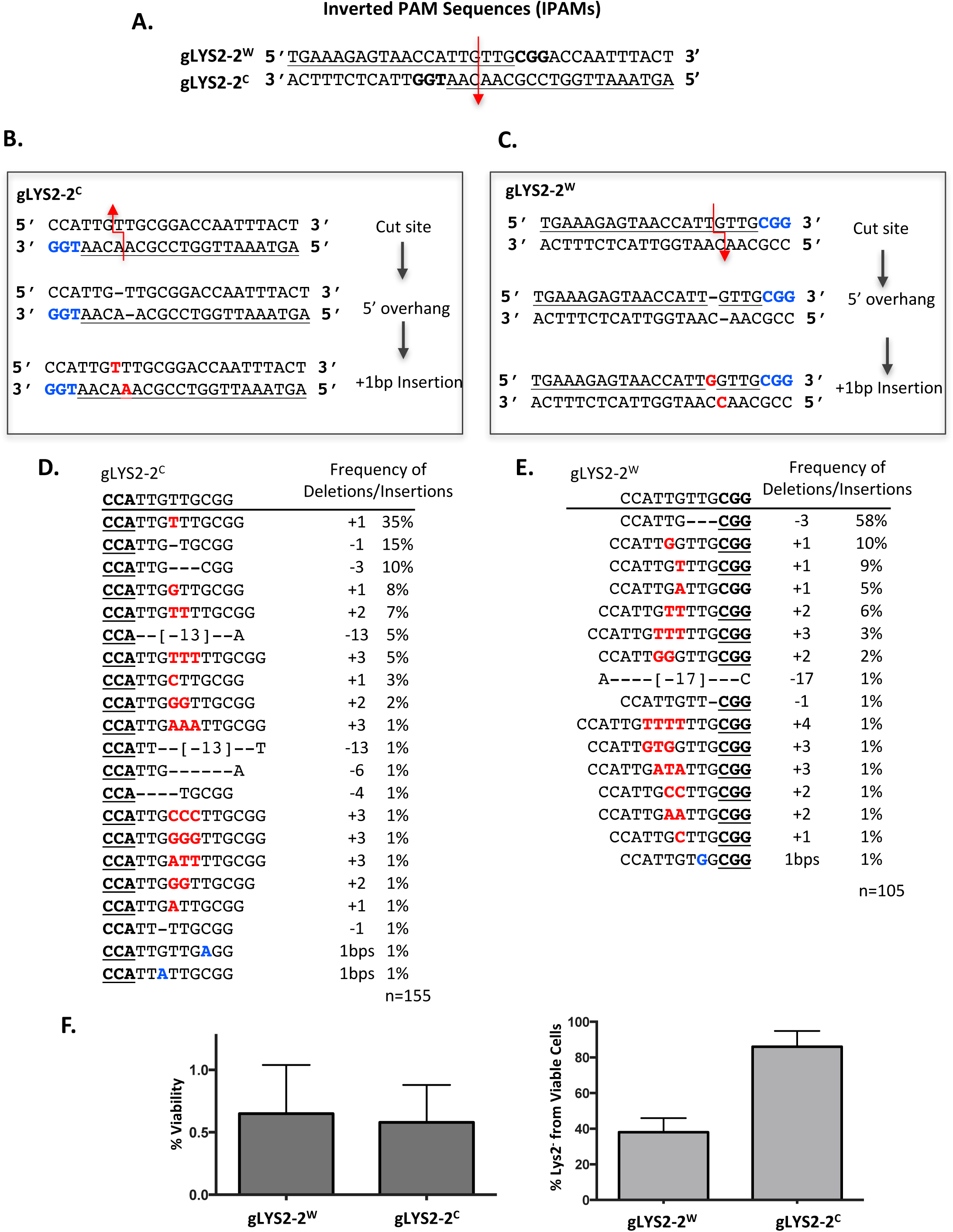
Templated insertions and nonrandom deletions at CRISPR/Cas9 DSBs. **A)** gLYS2-2^W^ and gLYS2-2^C^ target PAM sequences that are 3-bp from each other and should cut in the same DNA location. Arrow indicates predicted cut-site. **B)** Example of templated 1bp insertion following Cas9 induced DSB from gRNA targeting the LYS2 locus (gLYS2-2^C^). **C)** Example of templated 1bp insertion following Cas9 induced DSB from gRNA targeting the LYS2 locus (gLYS2-2^W^). **D)** Sequence profile survivors after gLYS2-2^C^ induced DSB. Most common event is a T insertion **E)** Sequence profile of survivors after gLYS2-2^W^ induced DSB. The most common event is a deletion of three bases (TTG), apparently by annealing between microhomologies. The second most common event is a templated 1bp insertion. **F)** *Left*: Viability of cells targeted with inverted PAM Sequences (iPAMs), gLYS2-2^W^ and gLYS2-2^C^. *Right*: Percentage of survivors that are Lys2-.

The observation that +1 insertions were not random, but apparently templated by the nature of the Cas9 cleavage, led us to create an ostensibly identical DSB by using a PAM on the opposite (Watson) strand directed by gLYS2-2^W^. For example, in this case the 5th base from the PAM would yield insertion of a G (reading the top strand) (Fig. 1C). We term the pair of PAMs as inverted PAM sequences (iPAMs) (Fig. 1A) and labeled the gRNAs as W or C, corresponding to a PAM located on the Watson (top) or Crick (bottom) strand. Overall, when cleavage is directed by the PAM on the “top” strand (gLYS2-2^w^), the outcomes were notably different from those directed to the opposite strand (Fig. 1D and 1E). Indeed, as predicted, the most prevalent +1 insertion was a G at the site predicted by a 1-nt 5’ overhang. However, there were nearly as frequent +1 insertions of T and A, although these were not at the predicted templated site (Fig. 1B). There were also a number of +2 and +3 insertions at the same site. In the case of gLYS2-2^C^, the most prevalent 2-base insertion (+TT) might arise also by a templated event if the cleavage left a 2-nt extensions, but templating cannot explain those for LYS2-2^W^. Indeed, here and in the data shown below, the +2 and +3 show a very strong bias toward homonucleotide insertions.

There were many other differences when comparing sequence profiles for the two gRNAs. For gLYS2-2^W^, about 40% of survivors were Lys2^-^ (Fig. 1F), whereas 80% of the outcomes were Lys2^-^ using the opposite gRNA. By far the most frequent event (58% of all events with LYS2-2^W^) was a 3-bp deletion involving the TTG sequences on either side of a blunt-ended DSB. Surprisingly, TTG microhomology-mediated deletions constituted only 9% for gLYS2-2^C^. There were other differences as well; a 13-bp deletion that was found 5% of the time for gLYS2-2^C^ was absent among the indels in gLYS2-2^W^ (Fig. 1C and 1E).

### +1 and other insertions mostly depend on Pol4

If there are 1-nt 5’ overhanging ends then there should be a DNA polymerase to fill in (5’ to 3’ on the recessed strand) before ligation could occur. We had previously implicated the PolX family DNA polymerase Pol4 in filling in misaligned DSB ends created by HO endonuclease (Moore and Haber, 1996). Here we show that most of the +1 templated events do indeed require Pol4 (Fig. 2). For both gLYS2-2^W^ and gLYS2-2^C^, the deletion of *POL4* eliminated nearly all the insertions – including the insertions of multiple bases (Fig. 2A). For gLYS2-2^W^, without Pol4, the frequency of 3-bp microhomology-bounded TTG deletions increased from 58% to 94% (Fig. 2C). With few to no insertions in the absence of Pol4, viability for both gRNAs decreased significantly (Fig. 2B). Additional evidence for the role of Pol4 in generating +1 and +2 insertions is presented for insertions created at another site (Supplemental Figure 1A).

**Figure 2.**
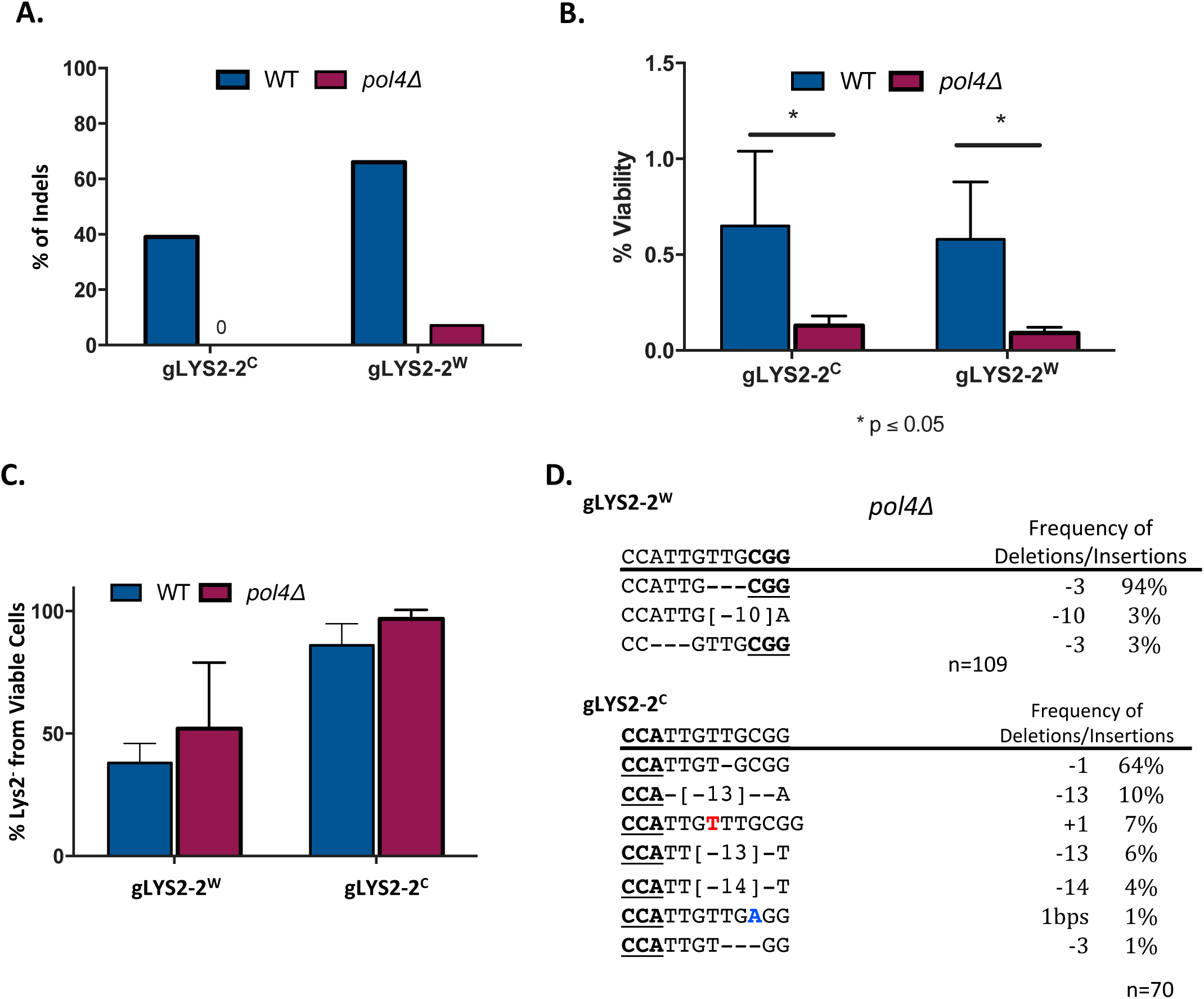
Pol4 is Required for Templated and Non-Templated Indels. **A)** Insertions during repair of gLYS2-2^W^ and gLYS2-2^C^ DSBs are mostly lost in the absence of *POL4*. **B)** Effect on NHEJ survival in the absence of *POL4*. **C)** Effect of *pol4Δ* on Lys2 mutations. D) Mutation profile of survivors following DSB from gLYS2-2^W^ and gLYS2-2^C^ in the absence of Pol4.

### Additional iPAMs sites confirm templated insertions and a strong bias in the overall NHEJ spectrum

To explore the templated nature of +1 insertions and the changes in the overall spectrum of indels depending on the orientation of Cas9, we examined 4 other iPAMs – three in other locations of the *LYS2* gene and one in *CAN1* (Figs. 3 and 4). gLYS2-3^C^ was inefficient in cleavage (Fig. 3A), but we could recover sufficient examples with an altered sequence to compare with gLYS2-3^W^ (Fig. 3D). The proportions of Lys2^+^ and Lys2^-^ cells with indels was statistically different for 2 of the 4 *LYS2* iPAMs (Fig. 3B). The two most common indels in each case are shown in Fig. 3D, with the complete distributions shown in Supplemental Table 1. For the iPAMs gLYS2-1^W^ and gLYS2-1^C^, deletions bounded by CT, TGTG and GAGA microhomologies predominated but their relative frequencies were noticeably different for the two gRNAs (Fig. 3D). As expected the most common +1 indel was a G for gLYS2-1^W^ and a T for gLYS2-1^C^ (Sup. Table 1). These trends are evident in the other pairs as well. For three of the gRNAs (gLYS2-3^W^, gLYS2-4^W^ and gLYS2-4^C^) many of the survivors had no indels, but instead had base pair substitutions that apparently rendered the gRNA ineffective. This is especially evident for gLYS2-4^C^, where the survival rate was less than 10% of the other gRNAs used. The low rate of viability cannot be attributed to an off-target cleavage by this gRNA, because viability proved to be nearly 100% when a 12-bp deletion of the protospacer – obtained from a survivor using gLYS2-4^W^ – was transformed with Cas9 and gLYS2-4^C^. The high rate of base-pair substitution is reminiscent of survivors of HO endonuclease cleavage when normal NHEJ was impaired and the rate of end-repair was only ∼5% of the wildtype rate (Moore and Haber, 1996). In addition to Sanger sequencing we also used Illumina sequencing® to sequence a larger pool of NHEJ survivors for gLYS2-4^C^ and gLYS2-4^W^. As shown in Supplemental Table 2, the proportion of mutations between the two sets of sequences for gLYS2-4^W^ were very similar. However, this was not the case for gLYS2-4^C^, where the Illumina sequencing data was much more varied than our previous Sanger sequencing results. Again, this difference can be attributed to the very low viability seen for gLYS2-4^C^ and the likely NHEJ-independent origin of many of the mutations recovered. Why Cas9 cleavage with gLYS2-4^C^ appears to block NHEJ whereas gLYS2-4^W^ does not, will require further investigation.

**Figure 3.**
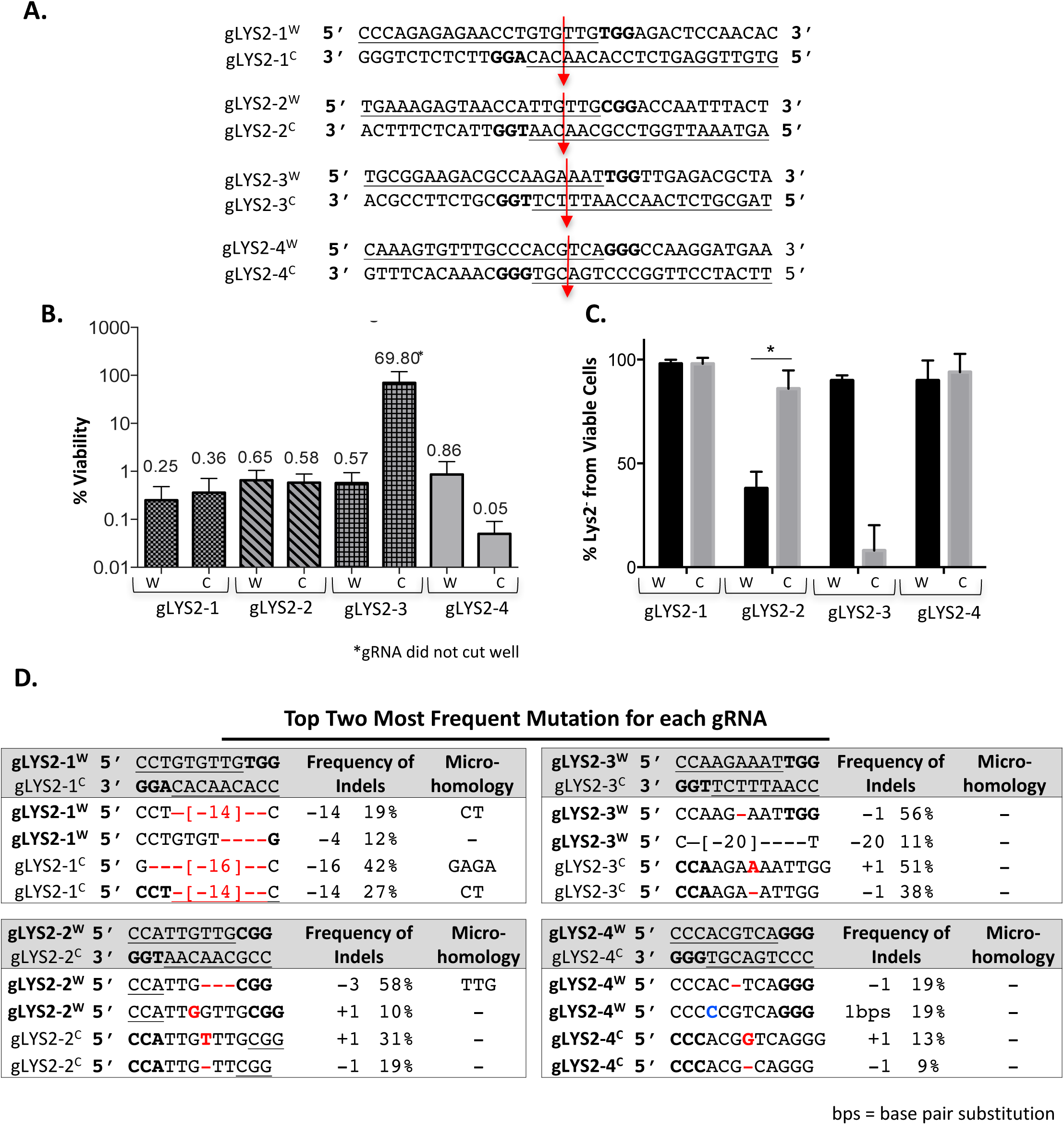
Inverted PAM sequences (IPAMs) have different indel repair profiles. **A)** Four pairs of gRNAs targeting the *LYS2* locus were designed (gLYS2-[1-4]) to cleave 4 different sites. Paired gRNAs were designed to create a DSB in the same DNA location, with gRNAs designated W or C, reflecting the location of the PAM. **B)** Percent viability from each gRNA targeting the *LYS2* locus. **C)** Percent of Lys2^-^ for each gRNA targeting *LYS2* from viable cells. A statistically significant difference in the percent Lys2^-^ cells was found for gRNA pair 2 (p =0.002). The very high proportion of Lys2^+^ in gLYS-3^C^ reflects the inefficient cutting with this gRNA; the great majority of Lys2^+^ had no change in the DNA sequence; hence no statistical comparison could be made with gLYS2-3^W^. **D)** Top two most frequent mutations from sequenced survivors for each gRNA. Many deletion events can be explained from microhomology mediated single-strand annealing.

Interestingly, we did not observe any of the predicted 1-bp templated insertions for gLYS2-3^W^ among 64 sequences. Instead, the most common event (56%) was a 1-bp deletion, which can be explained by resection of the ends and misalignment of an A-T base pair flanking the cleavage site (Supplemental Figure 3). This particular deletion could arise equivalently if the cleavage was blunt or with a 5’ overhang (Supplemental Figure 3A). In the case of gLYS2-2^C^, however, the prominent -A deletion (19% of survivors) can readily be explained if the cleavage had a 5’ overhang; however an analogous process beginning with a blunt end instead could produce a -G, which was not seen (Supplemental Figure 3A).

As noted above, at these other iPAM sites we also find insertions of 2, 3 or 4 nucleotides, which are almost always repetitions of the same base, and usually repetitions of the base that is templated in +1 insertions. There was one instance (gLYS2-1^W^) of a GTGGGGTGTGGG sequence that is reminiscent of telomere sequences (Sup. Table 1). Such captures of telomere sequences at DSB sites have been previously noted (Marzec et al., 2015).

A large proportion of the most frequent deletions we obtained are readily explained by annealing between microhomologies of 1-4 bases. In some cases (e.g. the 14-bp deletion bounded by CT microhomologies in gLYS2-1^W^and gLYS2-1^C^, Fig. 3D) there are comparable frequencies of the event in the two orientations. But among these same sequences, a frequent 5-bp deletion bounded by TGTG is absent with gLYS2-1^C^. Similarly, a 16-bp deletion, bounded by GAGA represents 42% of the sequences in gLYS2-1^C^ but only 4% of gLYS2-1^W^ (Sup. Table 1). These differences suggest that the position of Cas9 relative to the cleavage site exerts a strong influence on the outcomes.

We obtained similar results using iPAMs within the *CAN1* gene, where nearly all survivors had an indel and were canavanine-resistant (Fig. 4B). gCAN1-1^C^ led to 70% +1 (A) templated insertions, with rare insertions of C or T. However, with gCAN1-1^W^, only 16% proved to be templated +1 (G) insertions. Surprisingly, the most frequent event (23%) was a +1 (A) non-templated insertion. There were also a substantial number of +1 (T) insertions, inserted after the second base from the PAM (Fig. 4C). As we discuss below, these differences may reflect some error-prone fill-in synthesis that is strongly context dependent. Again, for gCAN1-1^W^ there were a number of insertions of 2, 3 and 4 bases that were mostly repeats of A, the most-frequently inserted, but apparently non-templated inserted base. Finally, we note a variety of deletions, some bounded by a single microhomology, which were not the same for the two gRNAs.

**Figure 4.**
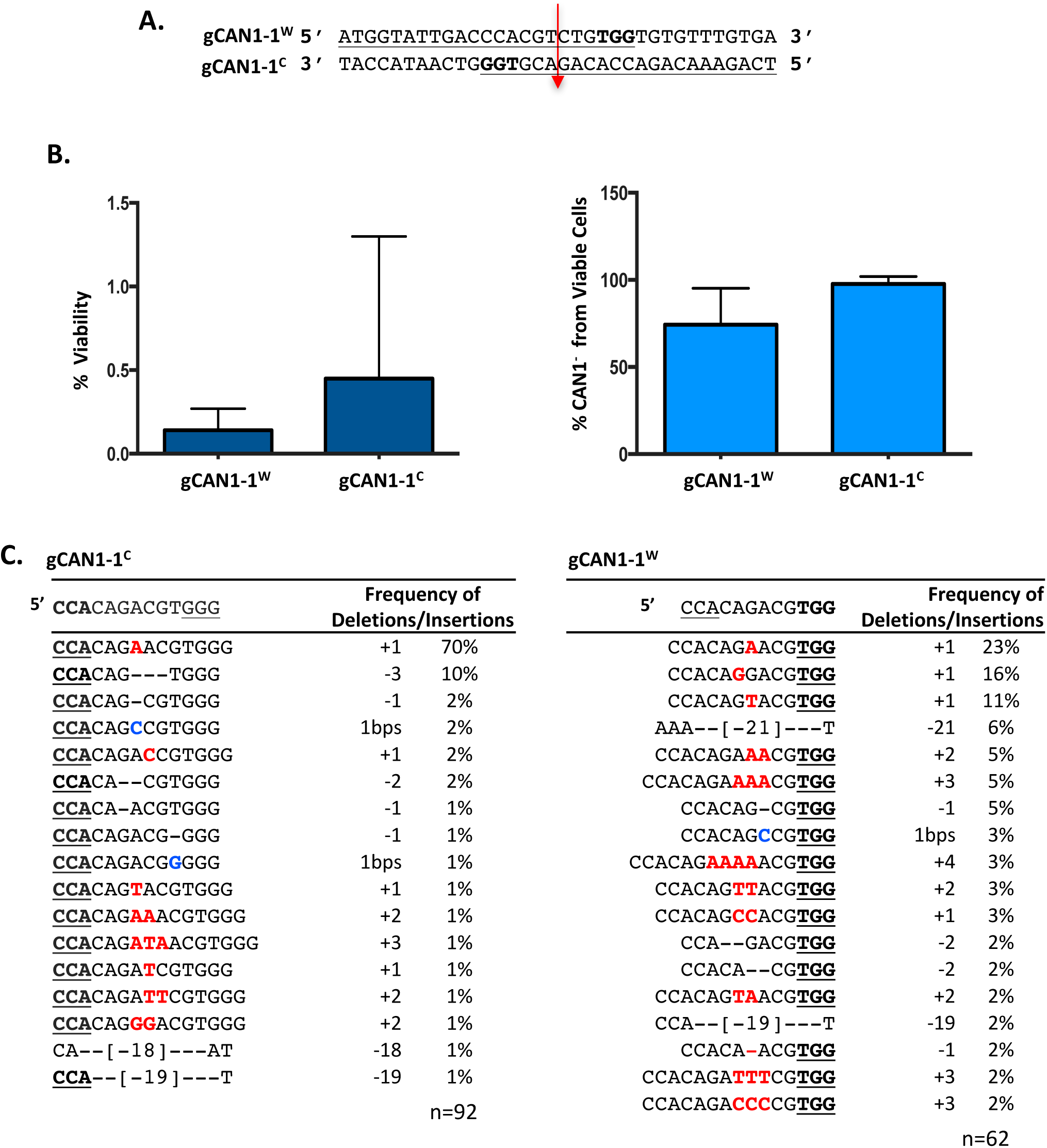
Templated Insertions after Cas9 induced DSB in *CAN1* Locus. **A)** gRNAs gCAN1-1^W^ and gCAN1-1^C^ target PAM sites on opposite strands of *CAN1* gene, but are expected to create the same DSB, 3-bps away from PAM site. **B)** Viability of cells following gCAN1-1^W^ and gCAN1-1^C^ expression. **C)** Indel profile from survivors from gCAN1-1^W^ and gCAN1-1^C^.

In Supplemental Figure 4 we also include results from a single gRNA in *CAN1*, showing again a strong bias towards templated +1 insertions (29%) as well as homonucleotide insertions (+AA, +AAA and even +AAAAA, as well as +TT and +TTTTT).

### Cas9-mediated indels are strongly dependent on the MRX complex

As previously reported, (Moore and Haber, 1996), indels within the HO cleavage site in *MATα*, even an in-frame 3-bp deletion or insertion, results in a *matα*1 mutation, causing sterility. We designed two gRNAs (*α*#1 and *α*#2) to cut in the same *MATα*1 gene, 23 and 96-bp away from the HO cleavage site (Fig. 5A). Cells were transformed with two plasmids, one containing a galactose-inducible Cas9 gene and the other expressing the appropriate gRNA. As expected, with continued Cas9 cleavage in the *MATα*1 gene, nearly all of the survivors were sterile, (*α*#1: 39/40, *α*#2: 38/40); the few non-sterile colonies proved to not have been mutated. The viability in GAL::Cas9 strains was approximately 5 times higher than with GAL::HO (Fig. 5B); this difference is likely a reflection of the slower kinetics of cleavage by Cas9 and the possibility that cells replicate before cleavage has occurred. As with HO DSBs, Cas9-mediated NHEJ strongly requires DNA ligase 4 and the Ku proteins, as *dnl4Δ* and *yku70Δ* derivatives displayed an approximately 100-fold decrease in viability (Fig. 5B).

**Figure 5.**
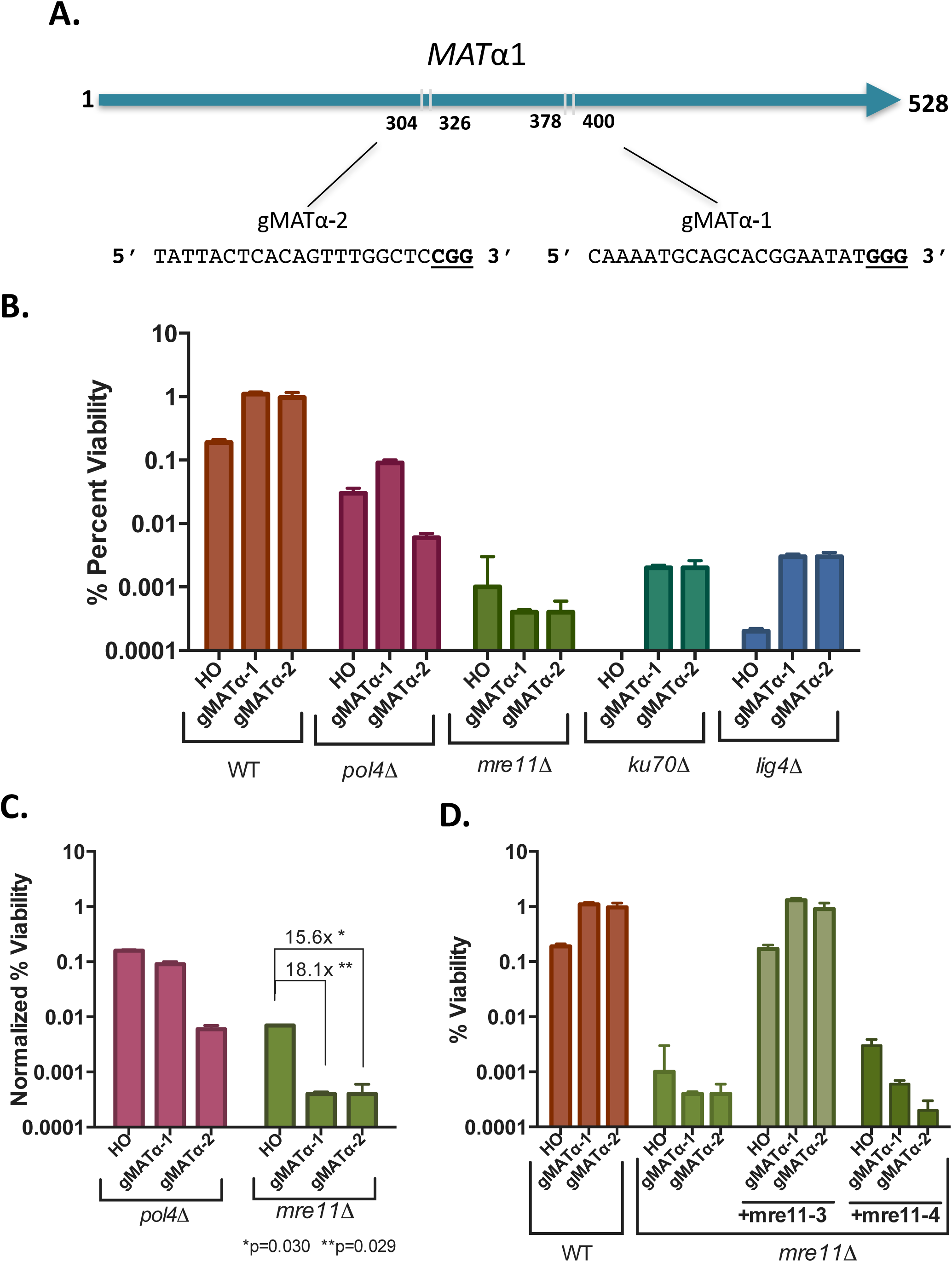
Mre11 is critical for Cas9 induced NHEJ Repair. **A)** Two gRNAs were used to target the *MATα* locus in the Yα sequence. **B)** Viability for HO, and CRISPR-Cas9 with g*MAT*α-1 or g*MAT*α-2 gRNAs in donorless strains that cannot repair the DSB by homologous recombination. **C)** Viability relative to WT for HO and CRISPR-Cas9 MAT g*MAT*α-1 or g*MAT*α-2 gRNA. Error bars represent standard error of the mean. **D)** NHEJ repair requires the MRX complex, independent of Mre11’s nuclease activity. Error bars represent standard error of the mean.

The Mre11-Rad50-Xrs2 (MRX) complex has been implicated in many steps of DSB repair, including the bridging of broken DNA ends (Bressan et al., 1999; Paull and Gellert, 2000; van den Bosch et al., 2003). With HO-induced DSBs, MRX proved to be important in the templated +2 and +3 insertions but was not required for a 3-bp deletion that also involved misaligned base-pairing of the overhanging ends (Moore and Haber, 1996). Here we examined the role of Mre11 in Cas9-induced NHEJ. Deletion of *MRE11* resulted in a more than 1000-fold reduction in survivors; moreover, among the *mre11 Δ* survivors only 8/367 had actually altered the cleavage site sequence; the rest were presumed to be rare cells that failed to induce cleavage of the target site, as we saw previously among rare survivors of HO cleavage (Moore and Haber, 1996). In these 8 cases, the altered target sequence either harbored a spontaneous mutation in the PAM or PAM-adjacent sequences (2 instances) or an indel, all of which were 1-bp deletions (Sup. Fig. 2). Thus, NHEJ of Cas9 cleaved DSBs was substantially more dependent on MRX than the HO breaks (Fig. 5C), which suggests that Mre11 is required not only for fill-in events but for nearly all deletions. This difference most likely reflects the fact that HO ends provide a 4-nt 3’ overhang to facilitate microhomology-mediated annealings whereas Cas9 cleavages are mostly blunt-ended and may therefore need the MRX complex either to initiate resection of the ends or to tether the ends together.

To test whether Mre11 nuclease activity or DNA end-bridging by the MRX complex was critical for Cas9 NHEJ repair, we investigated the repair efficiency in *mre11*Δ strains co-transformed with centromere-containing plasmids expressing either the nuclease-dead *mre11-3* allele or the *mre11-4* allele which fails to form a MRX complex (Bressan et al., 1999), transcribed from their native promoter. *Mre11-3* rescued the strain viability, whereas the *mre11-4* allele did not (Fig.5D). Importantly, the *mre11-3* allele rescued *mre11*Δ NHEJ viability for HO induction as well (Fig. 5D). These results suggest that Mre11’s nuclease activity is not required for its role in NHEJ of Cas9-generated ends but the integrity of the MRX complex is needed to allow efficient end-joining.

## DISCUSSION

CRISPR-Cas9 has democratized easily accessible genetic engineering, and therefore made it possible to create mutations and gene editing in many organisms that are not as genetically amenable to study as is *S. cerevisiae* (Jasin and Haber, 2016). Nonetheless, budding yeast still provides an important platform to analyze the mechanisms of Cas9-mediated events in great detail especially considering the potential applications of Cas9 in organism engineering. Here, we have studied NHEJ repair profiles in response to Cas9-mediated DBS’s and have reached several important conclusions.

First, we show that Cas9 induces a significant fraction of +1 indels that are apparently templated from a 1-bp 5’ overhang at the Cas9 cleavage site. What proportion of Cas9 cleavages are not blunt-ended is not known; *in vitro* the great majority of ends are certainly blunt (Jinek et al., 2012). However, *in vivo*, a 1-bp 5’ extension may facilitate recovery after NHEJ events, possibly by more efficient recruitment of an end-joining factor (Liang et al., 2016). In addition, an all-atom dynamics simulation predicted that Cas9 DNA cleavage could have 1-bp 5’ overhangs (Zuo and Liu, 2016). In budding yeast, blunt end ligations, as measured by plasmid ligation after transformation, are markedly less efficient than those with complementary overhanging ends (Daley et al., 2005). It is not yet known how efficiently Cas9-generated blunt ends are ligated *in vivo*; such a measurement requires that – after cleavage – Cas9 is rendered inactive to prevent re-cleavage of the perfectly joined ends. Experiments are underway in our lab to achieve the necessary synchronous induction and subsequent inactivation of cleavage that is required for such a measurement. We note again that finding +1 insertions is not restricted to budding yeast, but is readily seen in the spectra of indels in mammalian cells (Canver et al., 2017; Veres et al., 2014; Wang et al., 2014). Moreover, the non-random end-joining events described for an extensive study of different gRNAs by van Overbeek et al. (2016) show a very strong bias for the +1 insertion to be templated by the 4^th^ base from the PAM. Indeed, among 157 examples of Cas9-induced indels described by van Overbeek et al. (2016), in which there were at least 1000 +1 events, on average 90.8% of the +1 insertions at the expected insertion site were the base predicted from the sequence, with a range of 45 to 100% of the insertions being the predicted base.

Second, the filling-in of 5’ overhangs to produce +1 insertions generally requires the X family DNA polymerase, Pol4. In budding yeast Pol4 has been shown to be required for gap-filling between misaligned 5’ overhanging ends, although – in contrast to gap-filling of misaligned 3’- overhangs which totally depend on Pol4 – other polymerases appear to be able to substitute for Pol4 at a reduced efficiency (Liang et al., 2016). The residual insertions we find without Pol4 may thus depend on other polymerases able to create a blunt end. Pol4 lacks proofreading activity and is thus error-prone (Bebenek et al., 2005). This may explain the appearance of other bases instead of the templated base in some junctions. The frequency of recovering alternative bases is quite different in different gRNA contexts.

In addition to +1 insertions, Pol4 appears to be required when there are multiple base insertions at the break. The origin of these 2, 3 and 4-base insertions in budding yeast might be explained by the fact that Pol4 belongs to the X family of DNA polymerases, some of which have terminal transferase activity (Li et al., 2012; Loc'h et al., 2016; Nick McElhinny et al., 2005). Surprisingly, these multiple-base insertions are overwhelmingly non-random, with >85% of them being homonucleotide runs, the majority of which match the base most frequently inserted as the templated +1 insertions. For example, for gLYS2-2^C^, the templated base is T and the most frequent 2 and 3 bp insertions are also TT and TTT. Less often, there were +1 G insertions and there were also GG and GGG inserts. These may reflect some sort of slippage at the 1-nt overhang or some sort of terminal transferase activity. Among the multiple-base homonucleotide insertions 26% were runs of A, 54% incorporated T, 12% were Gs and 8% were C. Some bias likely reflects which iPAM sequences were chosen. Among the 5 iPAMs we analyzed, there were no predicted +1C insertions. It is also possible that Pol4 could copy the gRNA (Wei Yang, personal communication).

Inspection of indel data from an extensive study in human cells (Jiang et al., 2014; Seeger and Sohn, 2016; van Overbeek et al., 2016) also revealed the presence of +2 insertions at the same site where +1s are created. Overall, +2 insertions appeared at a frequency of approximately 8% of the number of +1 insertions at the same site. We examined 82 instances in which there were 200 or more +2 insertions. Among this set, when there were +1 insertions, the most frequent insertion was the expected templated event in all but 6 cases. Greater than 90% of all +1 insertions appear to be templated. Overall, the +2 inserts began with the same base as seen in +1 cases 85% of the time (ranging from 30% to 100%). We do not see evidence of homonucleotide +2 insertions as seems to be the case in budding yeast; however, in 57 of these 82 cases the most frequent +2 insertion can be accounted for if it, too, was templated, ostensibly by a +2 5’ overhang. Indeed, 76% of all insertions were the predicted +2 insert in these 57 cases. Among the 22 cases where the most frequent dinucleotide insertion was not that expected from a 2-nt 5’ overhang, the first base inserted was the expected base in all but 8 cases. Whether the additions of one or two bases will require PolX will be of great interest to determine.

Third, the use of palindromic PAM sequences allowed us to demonstrate clearly that the orientation of Cas9 on DNA strongly influences the spectrum of indels. Most striking is the case of gLYS2-2^W^ and gLYS2-2^C^ directed against *LYS2*. With a PAM on the non-transcribed strand >50% of the outcomes had a 3-bp deletion that would result from annealing TTG sequences on either side of the (blunt and resected) DSB, and another 10% were +1 templated insertions of a G (Fig. 1G). In contrast with a PAM on the transcribed strand, only 10% had the 3-bp deletion and 30% were the expected templated +1 insertion (+T) (Fig. 1C). These differences in outcome could reflect *in vitro* studies showing that Cas9 remains tightly bound to the site of cleavage, but releasing the end further from the PAM (Richardson et al., 2016). A large difference in the microhomology-mediated annealing of the flanking TTG sequences would also be expected if one of the two orientations led to a much larger proportion of 5’ overhanging cuts. Whereas blunt end cleavage would lead to both ends having TTG on the top strand, allowing perfect annealing of the termini after resection, cleavages producing 1-bp 5’ overhanging (resulting in TT and GTTG on the top strand) would not have precisely complementary microhomologies at the two ends. Whether there is an influence of the orientation of Cas9 relative to transcription also needs to be investigated systematically, but from the 5 iPAMs that we investigated there does not seem to be a clear pattern that one orientation or the other is more efficient in cleavage *per se* or in generating indels after cleavage (Fig. 3).

Finally, NHEJ events after Cas9-mediated cleavage are nearly completely dependent on the MRX complex. Two separation-of-function mutants have led us to the conclusion that NHEJ does not depend on the nuclease activity of MRX but does depend on the integrity of the complex. We suggest that the end-tethering promoted by the MRX complex becomes essential when the DSB ends are blunt. We note that NHEJ of Cas9 cleavages are not more sensitive either to the absence of Ku or DNA ligase 4.

Consistent with other studies of Cas9-mediated indels, we found that most of the deletions we recovered are mediated by microhomology present near the break site (Bae et al., 2014; Li et al., 2015; Nakade et al., 2014; Zhang and Matlashewski, 2015). Keeping this in mind may be helpful when designing gRNAs for NHEJ-induced loss of function. For example, almost 60% of cells expressing gLYS2-2^C^ targeting *LYS2* retained protein function and still grew on Lys2 dropout plates, this was due to a 3-bp deletion resulting from 3bp microhomology, maintaining the protein in-frame (Fig. 1D and 1G).

In conclusion, we find that the types of indels recovered by a particular gRNA-directed cleavage is strongly dependent on DNA sequence context, even if two PAMs are selected that direct cleavage to the same target. The basis of these biases remains an important subject to investigate.

## Acknowledgments

We are grateful to James DiCarlo and George Church for providing the yeast Cas9 expression plasmid. Research was supported by NIH grants GM20056 and GM76020 to J.E.H. B.L., A.K., and D.P.W. were supported by NIH Genetics training Grant T32 GM007122. J.E.B. was supported by the Brandeis University Provost’s Undergraduate Research Fund.

## Figure legends

**Supplementary Figure 1. NHEJ events for CRISPR Cas9 gRNAs g*MATα*-1, g*MATα*-2 gRNA g*MAT*a-1 and g*MAT*a-2 gRNA in WT and *pol4Δ* background A)** DNA sequences from cells targeted with either gRNA g*MATα*-1 or g*MATα*-2 in WT and *pol4Δ* background. Insertions are shown in red and deletions are represented in dashes. The PAM sequence is noted in blue. Number of bases inserted and or deleted indicated in brackets, with the frequency of each event to the left. **B)** g*MAT***a**-1 or g*MAT***a**-2 gRNA strand targeting sequences and mutation profiles in WT and *pol4Δ* background. **C)** Histogram of insertions/deletions for each gRNA in WT and *pol4Δ* background.

**Supplementary Figure 2.** Cas9 NHEJ events for *mre11*Δ, *mre11*Δ + *mre11-3* CEN, and *mre11Δ* + *mre11-4* CEN strains. Insertions shown in red lettering, whereas deleted bases are marked by dashes. PAM sequences indicated in blue and underlined. Number of bases inserted and or deleted indicated in brackets, with the frequency of each event to the left of each indel.

**Supplementary Figure 3. 5’ overhangs can also account for -1 deletions in some instances.** Possible origins of a -1 deletion are compared for blunt or 1-nt 5’ overhangs for two examples. In A, resection and annealing of a single base, followed by clipping of an unpaired base and filling-in any gaps, can result in a -1 deletion whether the DSB is blunt or overhanging. In B, however, the blunt end deletion yields a -1 deletion that is not observed (-1G), whereas the 5’ overhang will provide a route to create the -1 A deletion that is frequently recovered.

**Supplementary Figure 4.** Cas9 NHEJ events in another site in *CAN1* show +1 templated insertions. The second most frequent event is a 7-bp deletion that can be explained by microhomology-mediated end-joining between adjacent TGG sequences.

**Supplementary Table 1.** Mutation profile from survivors of iPAMs gRNAs targeting *LYS2.*

**Supplementary Table 2.** MiSeq mutation profiles from ∼1000 survivors of iPAM gLYS2-4^C^ and gLYS2-4^W^.

**Supplementary Table 3.** Sequences of gRNAs used in this paper, shown as their DNA equivalent.

